# Brain virtual histology with X-ray phase-contrast tomography Part II: 3D morphologies of amyloid-β plaques in Alzheimer’s disease models

**DOI:** 10.1101/2021.03.25.436908

**Authors:** Matthieu Chourrout, Margaux Roux, Carlie Boisvert, Coralie Gislard, David Legland, Ignacio Arganda-Carreras, Cécile Olivier, Françoise Peyrin, Hervé Boutin, Nicolas Rama, Thierry Baron, David Meyronet, Emmanuel Brun, Hugo Rositi, Marlène Wiart, Fabien Chauveau

**Author notes:** Faculty of Medicine, The Ottawa Hospital and University of Ottawa, Ottawa, Ontario, Canada (Carlie Boisvert). Co-last authors (Marlène Wiart, Fabien Chauveau). Corresponding author: Fabien CHAUVEAU, Lyon Neuroscience Research Center, Groupement Hospitalier Est, Bât. CERMEP-Imagerie du Vivant, 59 BOULEVARD PINEL, 69677 BRON cedex FRANCE, Mail. **Author contribution** (according to Contributor Role Taxonomy (CRediT)): Conceptualization: MW, FC, Data curation: MC, MR, CG, CO, EB, HR, Formal analysis: MC, MR, Funding acquisition: FP, HR, MW, FC, Investigation: MC, MR, CB, CG, CO, Methodology: EB, HR, MW, FC, Project administration: MW, FC, Resources: DL, IAC, HB, NR, TB, DM, Supervision: EB, HR, MW, FC, Validation: MC, FC, Visualization: MC, MR, CB, CG, Writing – original draft: FC, Writing – review & editing: MC, MR, CB, EB, HR, MW, FC.

## Abstract

While numerous transgenic mouse strains have been produced to model the formation of amyloid-β (Aβ) plaques in the brain, efficient methods for whole-brain 3D analysis of Aβ deposits are lacking. Moreover, standard immunohistochemistry performed on brain slices precludes any shape analysis of Aβ plaques. The present study shows how in-line (propagation-based) X-ray phase-contrast tomography (XPCT) combined with ethanol-induced brain sample dehydration enables hippocampus-wide detection and morphometric analysis of Aβ plaques. Performed in three distinct Alzheimer mouse strains, the proposed workflow identified differences in signal intensity and 3D shape parameters: 3xTg displayed a different type of Aβ plaques, with a larger volume and area, greater elongation, flatness and mean breadth, and more intense average signal than J20 and APP/PS1. As a label-free non-destructive technique, XPCT can be combined with standard immunohistochemistry. XPCT virtual histology could thus become instrumental in quantifying the 3D spreading and the morphological impact of seeding when studying prion-like properties of Aβ aggregates in animal models of Alzheimer’s disease. This is Part II of a series of two articles reporting the value of in-line XPCT for virtual histology of the brain; Part I shows how in-line XPCT enables 3D myelin mapping in the whole rodent brain and in human autopsy brain tissue.

**Highlights:**

- X-ray phase-contrast tomography (XPCT) enables whole brain detection of Aβ plaques
- Morphometric parameters of Aβ plaques may be readily retrieved from XPCT data
- New shape parameters were successfully extracted from three Alzheimer’s disease models
- A Fiji-based “biologist-friendly” analysis workflow is proposed and shared
- XPCT is a powerful virtual histology tool that requires minimal sample preparation

Graphical abstract

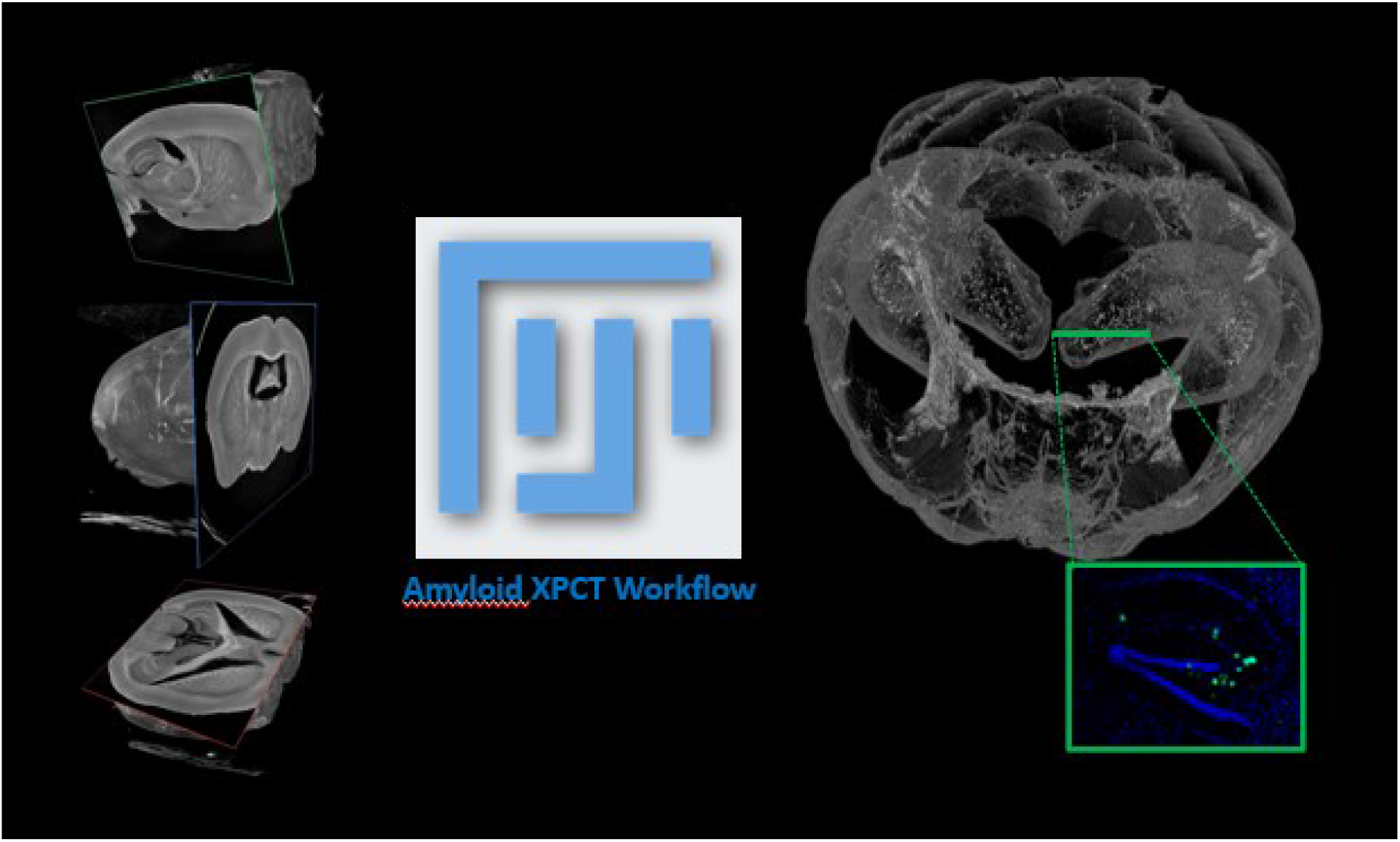

## Introduction

The amyloid cascade is considered pivotal in the development of Alzheimer’s disease. In the last 20 years, numerous transgenic mouse strains have been produced to model the accumulation of amyloid-β (Aβ) peptides and formation of Aβ plaques in the brain. Most are knock-in animals, obtained by insertion of human genes, which usually include mutations observed in familial cases: presenilins (PSEN1/2), amyloid precursor protein (APP), microtubule associated protein tau (MAPT), triggering receptor expressed on myeloid cells 2 (TREM2), etc. The Alzforum database currently lists 123 mouse models with 1 transgene, and 62 multi-transgene mouse models (https://www.alzforum.org/research-models; last access on May 25, 2021). Fast methods for whole-brain analysis are thus of great value for characterizing and comparing phenotypes among the variety of strains available.

X-ray based virtual histology is a new field of research mainly aimed at 3D datasets of biological tissue virtually sliced in any direction [1]. X-ray phase-contrast tomography (XPCT) using synchrotron radiation allows imaging of excised biological tissues or organs with weak X-ray absorption (like the brain) at microscopic level [2]. XPCT achieves a high signal-to-noise ratio without the need to add staining agents, by probing small changes in refractive indices in the tissue microstructure. It provides micrometric spatial resolution and isotropic reconstruction in a ~ cm^3^ field of view (FOV), thus offering ideal prerequisites for imaging protein aggregates of ~ 10μm in the entire and intact (unsliced, unstained) mouse brain [3].

Several pioneer studies already reported detection of Aβ plaques with various X-ray phase contrast techniques [4–6]. However, these first developments required long acquisition times (30-180 min) and/or provided limited anatomical contrast. Recently, free space propagation between object and detector has been shown to be a simple, fast and efficient technique for detecting Aβ plaques with good contrast [7,8]. In addition, anatomic contrast can be greatly enhanced by dehydrating the brain prior to imaging, which increases small local differences between the refractive indices of the different brain structures [2,3]. In terms of analysis and quantification, most previous studies reported only qualitative results, or quantification restricted to parameters accessible to standard 2D histology (e.g., number of plaques, diameter, volume). Only one recent report used XPCT to extract sphericity of Aβ plaques inside the cerebellum [9]. The present study aimed to realize the full potential of XPCT by combining:

i. optimal brain tissue preparation through dehydration in ethanol,
ii. fast acquisition in three different (mono, double and triple) transgenic mouse strains displaying Aβ pathology,
iii. semi-automatic segmentation of Aβ plaques inside the hippocampus using open-source tools (combination of Fiji plugins),
iv. and full characterization of their 3D morphology.

This is Part II of a series of two articles reporting the value of in-line XPCT for virtual histology of the brain; Part I shows how in-line XPCT enables myelin mapping of the whole brain [10].

## Methods

### Samples

All experimental procedures were carried out in accordance with European regulations for animal use (EEC Council Directive 2010/63/EU). The present study was performed on excised brains. Three transgenic lines were used, for a total of 8 brains:

i. mono-transgene line J20 (n=2 animals, 2 y.o.), with mutant APP [11];
ii. double-transgene line APPswe/PSEN1dE9 or APP/PS1 (n=3 animals, 1 y.o.), with mutant APP and mutant PSEN1 [12];
iii. triple-transgene line 3xTg (n=3 animals, 1 y.o.), with mutant APP, mutant PSEN1 and mutant MAPT [13].

One additional brain, from a wild-type mouse of C57BL6/129sv background, was used as a control.

### Preparation

Formaldehyde-fixed brains were dehydrated in ethanol and conditioned in plastic tubes (1 cm diameter) filled with ethanol.

### Acquisition

Imaging was performed at the ID19 beamline of the European Synchrotron Research Facility (ESRF, Grenoble, France). Acquisition parameters and data characteristics are summarized in Table 1. Briefly, the tomographic images were recorded within 3 min at a single sample-detector distance where the camera was positioned away from the sample (3 m) to obtain phase contrast. The experiments were performed with a polychromatic “pink” incident X-ray beam of 26 keV energy. Tomographic reconstructions were performed using the single distance phase-retrieval approach (“Paganin” method [14]) with PyHST2 software [15], the δ/β ratio being set to 1,000.

**Table 1-.** Acquisition parameters and data characteristics.

|  |  |
| --- | --- |
| **Detector** | sCMOS (PCO edge) |
| **Scintillator** | LuAg 500 $\mu\text{m}$ |
| **Energy [keV]** | 26 |
| **Sample-detector distance [m]** | 3 |
| **Exposure time [s]** | 0.04 |
| **Number of projections** | 3,500 – 3,600 (360°) |
| **Acquisition time [min]** | 3 |
| **Reconstruction algorithm ($\delta/\beta$)** | Paganin (1000) |
| **Field-of-view [$\text{cm}^3$]** | $\sim 1.3 \times 1.3 \times 0.9$ |
| **Matrix** | 2,048 x 2,048 |
| **Total slice number** | 1,400 |
| **Voxel size (isotropic) [$\mu\text{m}$]** | 6.5 |
| **Image size (type)** | 11.7 Go (16-bit) |

### Segmentation

Semi-automated detection of Aβ plaques used Fiji software [16]. A pipeline was built with the following plugins: segmentation editor (to isolate hippocampus: https://imagej.net/Segmentation_Editor), trainable WEKA segmentation 3D (to identify plaques) [17], and MorpholibJ [18] and 3D ImageJ suite [19] (to label objects and extract relevant parameters). Image features used for trainable segmentation were “Difference of Gaussian”, “Variance”, and “Maximum” for J20 and 3xTg. For APP/PS1, “Minimum” was used instead of “Maximum” after the application of a 3D Laplacian-of-Gaussian (Mexican Hat) filter [20], as suggested by Astolfo et al. [7]. A step-by-step guide to perform all image processing, “Amyloid-β XPCT Workflow”, is publicly available (DOI: 10.5281/zenodo.4584752) [21].

### 3D parameters

Sphericity, elongation, flatness, sparseness and distance to nearest object were outputs from 3D ImageJ Suite. Surface area, volume, mean breadth and mean signal intensity (here normalized by background intensity) were outputs from MorpholibJ. Definitions of these parameters are reported in Table 2.

**Table 2-.** Morphometric parameters. (More details are available at: https://imagej.net/MorphoLibJ.html#Documentation)

| Name | Definition | Formula |
| --- | --- | --- |
| **Compactness**<br>(not reported) | Ratio ($C$) between volume ($V$) and surface area ($A$); “3D ImageJ Suite” implements a ‘discrete’ compactness, described in [22] estimated from the number of voxels ($N_{\text{voxels}}$); spheres have a compactness of 1. | $C = \frac{36 \cdot \pi \cdot V^2}{A^3}$ $\approx \frac{N_{\text{voxels}} - \frac{A}{6}}{N_{\text{voxels}} - \left(\sqrt[3]{N_{\text{voxels}}}\right)^2}$ |
| **Sphericity** | Ratio ($S$) between volume ($V$) and surface area ($A$); linked to compactness (or discrete compactness); spheres have a sphericity of 1. | $S = C^{1/3}$ $\approx \left( \frac{N_{\text{voxels}} - \frac{A}{6}}{N_{\text{voxels}} - \left(\sqrt[3]{N_{\text{voxels}}}\right)^2} \right)^{1/3}$ |
| **Elongation** | Elongation of the equivalent ellipsoid, where $R_1 \geq R_2 \geq R_3$ are the lengths of the semi-axes of the ellipsoid with same inertia tensor as the object | $\text{Elongation} = \frac{R_1}{R_2}$ |
| **Flatness** | Flatness of the equivalent ellipsoid, where $R_1 \geq R_2 \geq R_3$ are the lengths of the semi-axes of the ellipsoid with same inertia tensor as the object | $\text{Flatness} = \frac{R_2}{R_3}$ |
| **Sparseness** | Ratio between the volume of the equivalent ellipsoid ($V_{\text{ell}}$) and the volume of the object ($V_{\text{obj}}$) | $\text{Sparseness} = \frac{V_{\text{ell}}}{V_{\text{obj}}}$ |
| **Mean Signal Intensity (SI)** | Mean intensity from the raw image; here reported as a mean signal-to-background ratio |  |
| **Volume** | Volume in the given unit ($\mu\text{m}^3$) | |
| **Surface Area** | Surface area in the given unit ($\mu\text{m}^2$) | |
| **Mean Breadth** | Average of the Feret diameter of convex objects measured over a selected number of orientations ($\mu\text{m}$) | |
| **Distance** | Distance between two closest segmented objects ($\mu\text{m}$) | |

### Analysis and statistics

Prism 8 (GraphPad) was used for violin plots and statistics. The Welch version of one-way ANOVA (gaussian populations, unequal variances) was used to compare the above-cited parameter values of individual Aβ plaques across the three transgenic groups, with additional Games-Howell tests (recommended for n>50) for pairwise comparisons. Linear correlations between parameters were quantified on Pearson correlation coefficient. MIPAV (v10.0.0, CIT, NIH [23]) was used for 2D visualization of semi-transparent overlay of plaque labels onto native images. Amira Software 6 (ThermoFisher) was used for all 3D renderings. *Immunohistochemistry*. One sample of each transgenic strain underwent paraffin embedding, 7μm-thick microtome slicing (Leica RM2245) and standard Aβ immunofluorescence or histological staining. Aβ immunofluorescence was performed after i) standard dewaxing, ii) antigen retrieval with 100% formic acid for 15-20 min at room temperature (RT), and iii) blocking with 5% Bovine Serum Albumin (BSA) and 0.5% Triton-X in Phosphate Buffer Saline (PBS, Sigma-Aldrich ref. P4417) for 30 min at RT, by incubating the anti-Aβ monoclonal antibody 4G8 (Covance, ref. SIG-39220, dilution 1:1,000 in 5% BSA) on each microscope slide overnight, in a humid chamber at 4°C. After three 5 min washes in PBS at RT, secondary antibody (anti-mouse AlexaFluor 488 or 546, ThermoFisher, dilution 1:500 in 5% BSA) was incubated on each microscope slide for 60 min, in a humid chamber at RT. After three 5 min washes in PBS at RT, slides were mounted with DAPI-containing medium (Roti-Mount FluorCare DAPI, Carl Roth), and stored at 4°C. Thioflavin S staining (ThS, Sigma-Aldrich ref. T1892) (0.05% in ethanol 70%) was performed for 10 min at RT after dewaxing, and slides were mounted with DAPI-containing medium, after two 1 min washes in ethanol 70%. Fluorescence images were captured on a microscope (AxioScope A1, Zeiss) equipped with a digital camera interfaced with image-analysis software (ZEN 2 lite, Zeiss).

### Data availability statement

The raw data required to reproduce these findings cannot be shared at this time due to the large size of the XPCT datafile, but are available on request. The image processing workflow is available on Zenodo (DOI: 10.5281/zenodo.4584752) [21].

## Results

### Qualitative analysis

After ethanol dehydration, brain anatomy, and specifically white matter tracts, was uniquely displayed on XPCT images (Fig. 1), as reported in Part I of this series of articles [10].

**Figure 1-.**
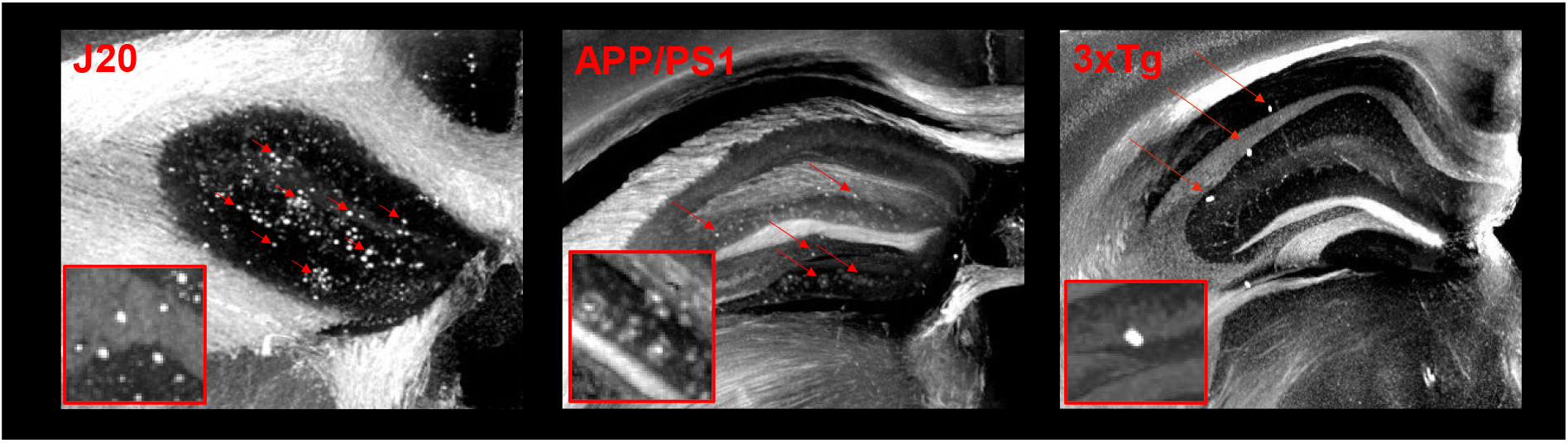
Whole-brain anatomy of ethanol-dehydrated brains from the three mouse strains (XPCT single slices).

One additional formaldehyde-fixed sample of the J20 strain was scanned in PBS using the same set-up and reconstruction algorithm (Fig. S1): though some Aβ plaques were visible, surrounding brain tissue exhibited low overall signal intensity, strongly contaminated by ring artefacts. Thus, imaging in ethanol was required for region-specific segmentation. In this proof-of-concept study, we chose to focus on the hippocampus, in which all three mouse strains exhibited Aβ plaques. However, these plaques displayed strikingly different appearances (Fig. 2): i) J20 exhibited numerous small intense spots, sometimes very close to each other and seemingly coalescing; ii) APP/PS1 displayed intense spots, often surrounded by a diffuse rim, resembling typical human dense-core plaques; iii) 3xTg showed few but large and highly intense deposits.

**Figure 2-.**
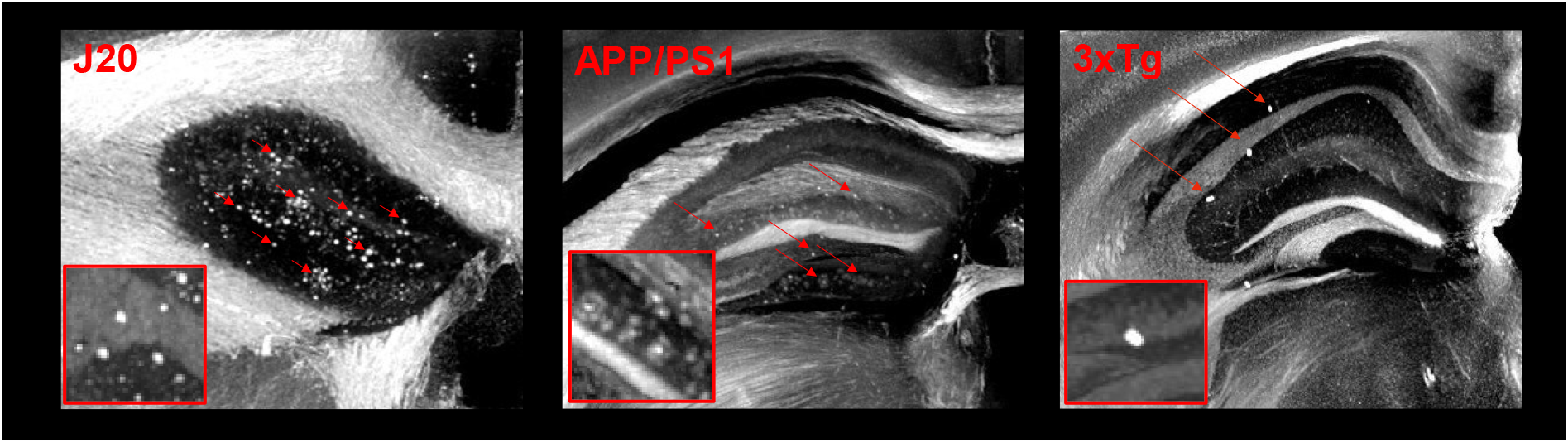
XPCT images (maximum intensity projections over 50 slices) obtained at the level of the dorsal hippocampus, showing the density and appearance of Aβ plaques in the three mouse strains.

These 3D signals obtained without labeling matched the 2D fluorescence of corresponding brain slices after thioflavin S staining or Aβ-peptide immunohistochemistry (Fig. 3), although some of the stained plaques seemed not to produce hyperintense contrast on XPCT.

**Figure 3-.**
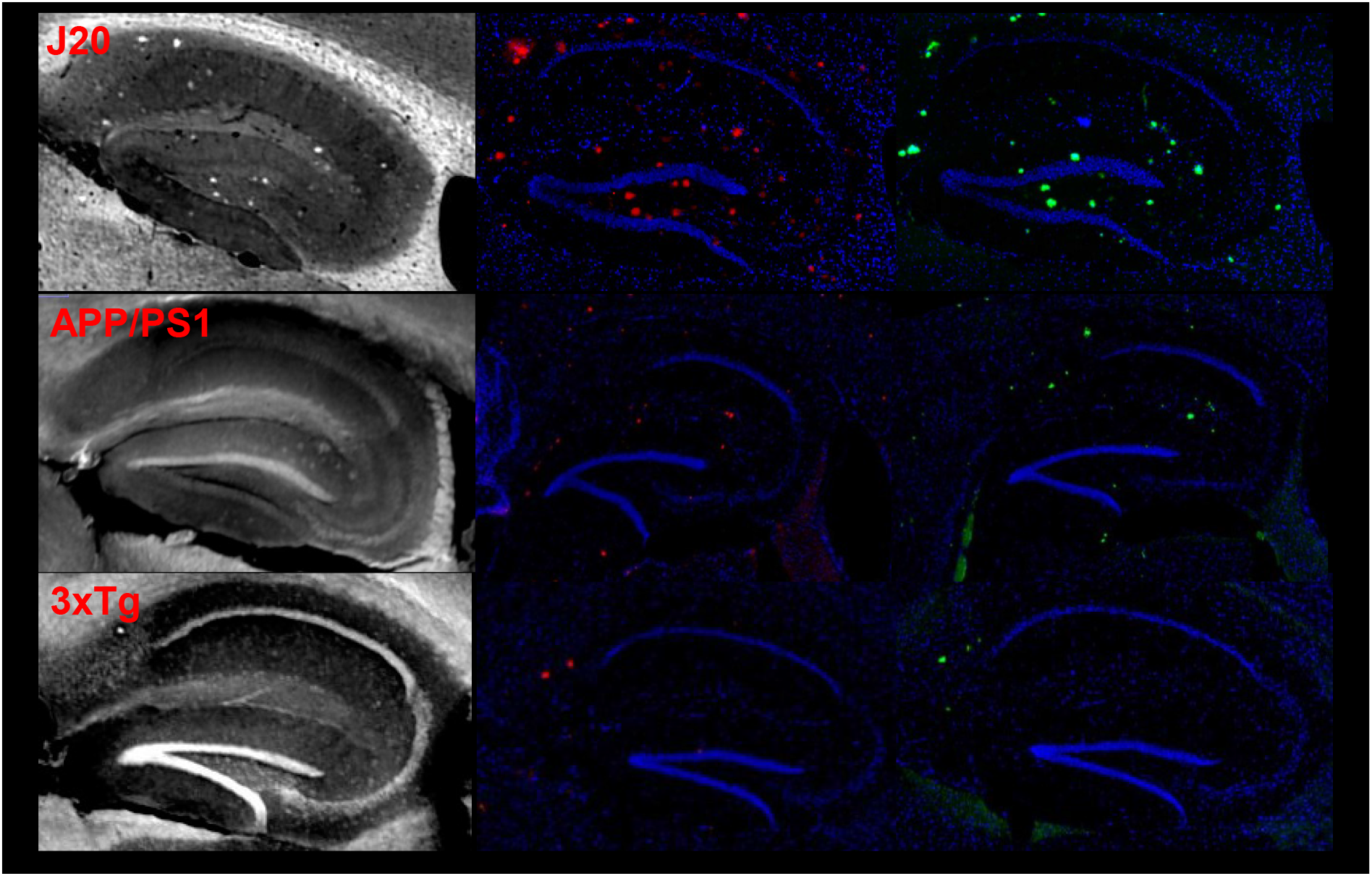
Corresponding XPCT image (left row), amyloid staining with 4G8 antibody (middle row, red), and Thioflavin S staining (right row, green), for the three transgenic mouse strains.

### Morphological quantification

Following hippocampus extraction (steps 2-3 in the Amyloid-β XPCT Workflow), machine learning was used to perform Aβ plaque recognition: trainable WEKA segmentation 3D used a strain-selective classifier that was built from 5 consecutive slices and then applied to the whole hippocampus (>200 slices) to produce a probability map that was thresholded by visual inspection (steps 4-5 in the Amyloid-β XPCT Workflow). This strategy proved versatile and accommodated the different types and numbers of plaques imaged in this study (Fig. 4).

**Figure 4-.**
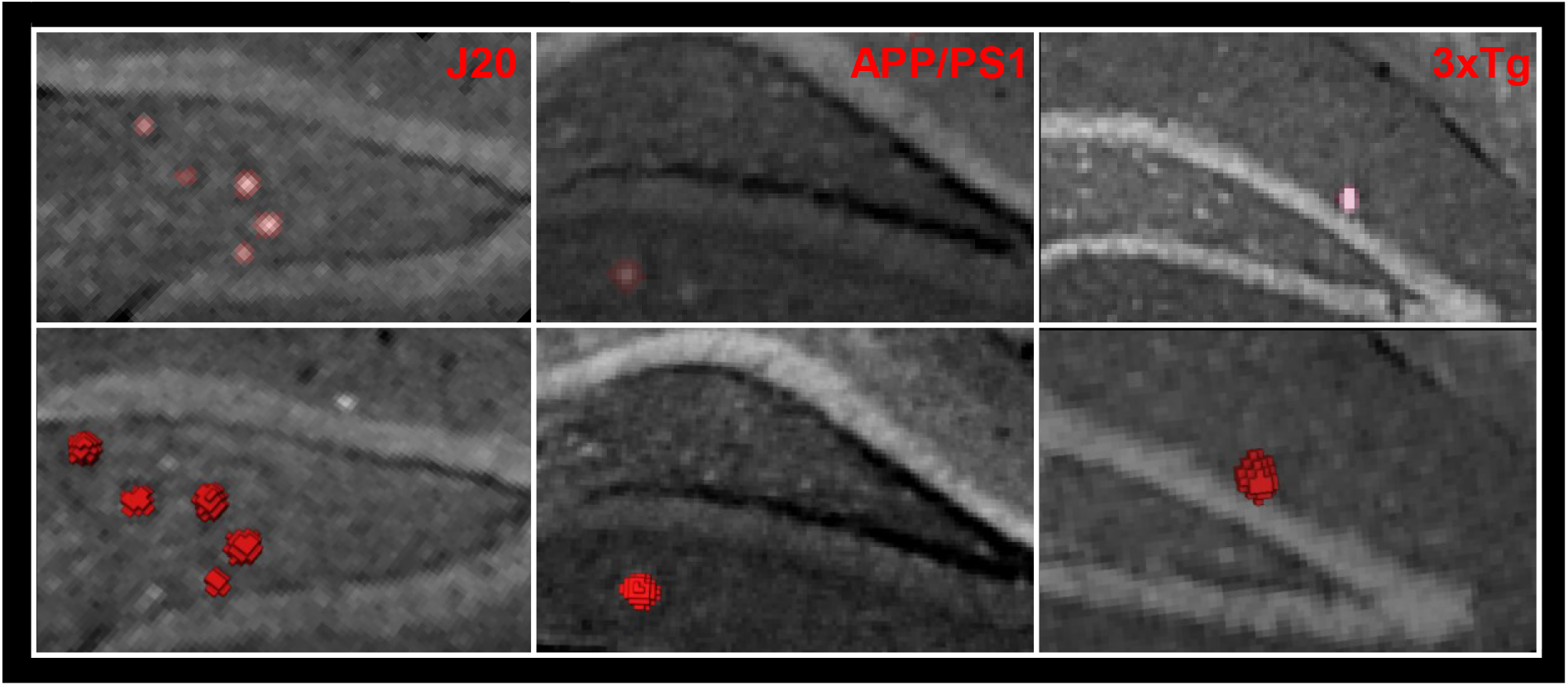
Representative 2D overlay and 3D rendering of segmented plaques for each strain.

After automatic labeling and size filtering of segmented objects (step 6 in the Amyloid-β XPCT Workflow), a few manual cleaning steps were performed under expert supervision (author FC). False-positive objects were, in most cases, located in the myelinated tracts of the perforant pathway (Fig. S2). On rare occasions, neurons or blood clots remaining in vessels were also detected as plaques. In 3xTg brains, a few false-positive labels were easily identified and removed, out of a dozen correctly segmented plaques (6 to 16 per hemisphere). In J20, which in contrast had several hundreds of plaques (774-916 per hemisphere), expert screening showed that the false-positive detection rate was below 5%. Importantly, not all J20 plaques could be individually separated; hence some segmented objects were “twin” plaques, in close contact, which were manually removed from analysis. In the case of APP/PS1, plaques were of lower signal intensity, and could not be unambiguously distinguished from perforant pathway signals. APP/PS1 volumes were thus screened to manually select a hundred representative plaques outside the perforant tracts, which were used to calculate morphological parameters. Finally, as a specificity control, the 3 classifiers were applied to the hippocampus of a wild-type mouse, and, after similar thresholding and size filtering, yielded no other detection than the same type of false-positive objects described above (Fig. S2).

Both the MorphoLibJ and 3D ImageJ Suite plugins were able to extract morphological parameters (step 7 in the Amyloid-β XPCT Workflow), and we here report a combination of the shape parameters available for each strain (Fig. 5).

**Figure 5-.**
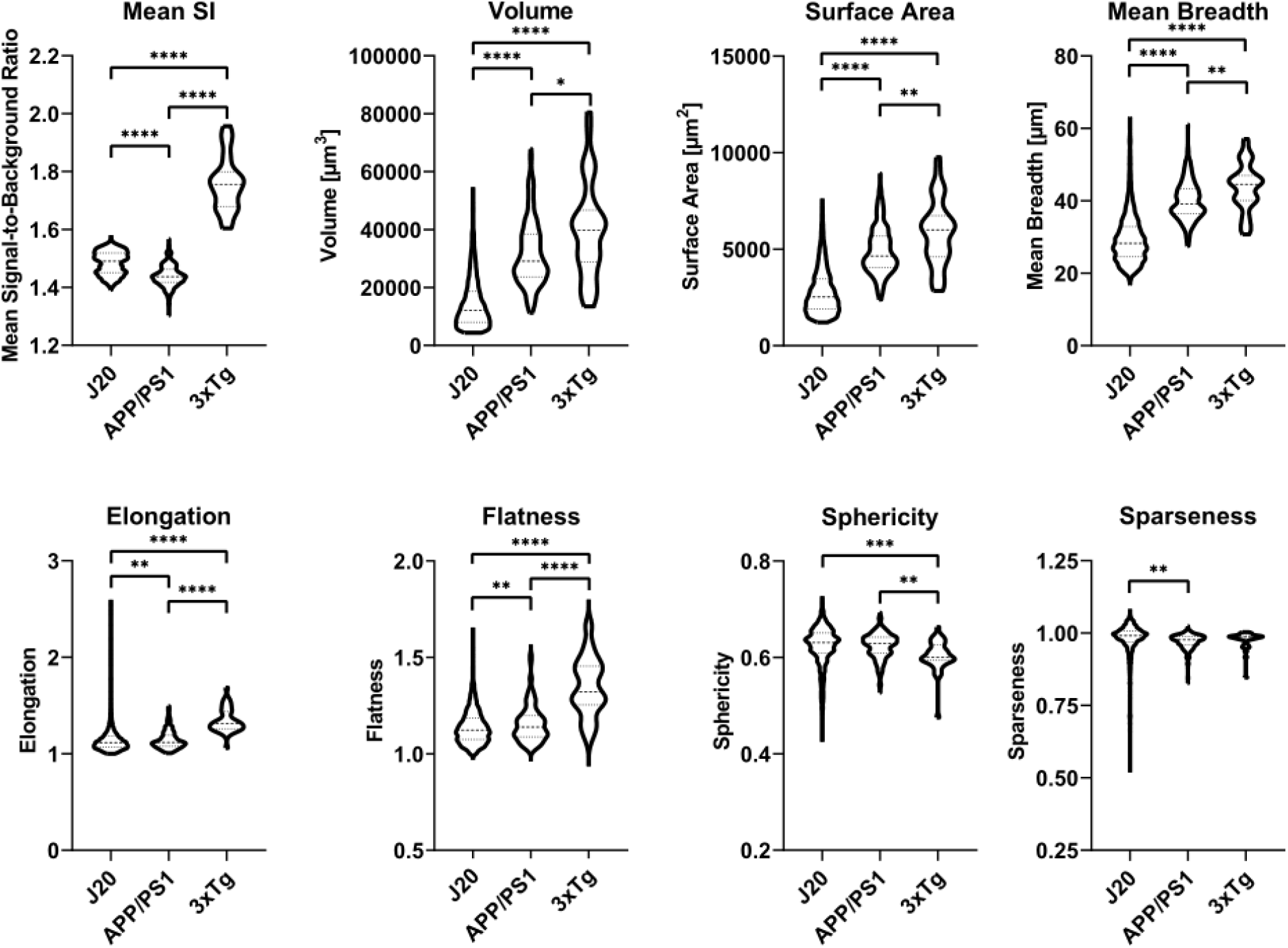
Violin plots of morphological parameters extracted from the population of Aβ plaques. Definitions of the parameters are reported in Table 2. Significance levels of pairwise Games-Howell comparisons are indicated (* p<0.05; ** p< 0.01; *** p<0.001; **** p<0.0001). Mean (interquartile range) are represented as dashed lines.

As expected, 3xTg displayed the highest plaque signal intensity after normalization to background (3xTg > J20 > APP/PS1, p<0.0001), while J20 had the smallest volume (J20 < APP/PS1 < 3xTg, p<0.0001) and surface area (J20 < APP/PS1 < 3xTg, p<0.0001) per plaque. Mean breadth, a parameter proportional to the integral of the mean curvature, was also significantly less in J20 (< APP/PS1 < 3xTg, p<0.0001). Shape differences were more pronounced for 3xTg, which had greater elongation and flatness (3xTg > APP/PS1 ≈ J20, both p<0.0001), and lower sphericity (3xTg < APP/PS1 ≈ J20, p=0.0002) compared to J20 and APP/PS1. Finally, it was possible to compute nearest-neighbor distances (Fig. S3), the mean value of which was 4-fold greater in 3xTg than J20.

These parameters are not mutually independent, and correlation matrices were computed to highlight strain differences in correlation coefficients (Fig. 6).

**Figure 6-.**
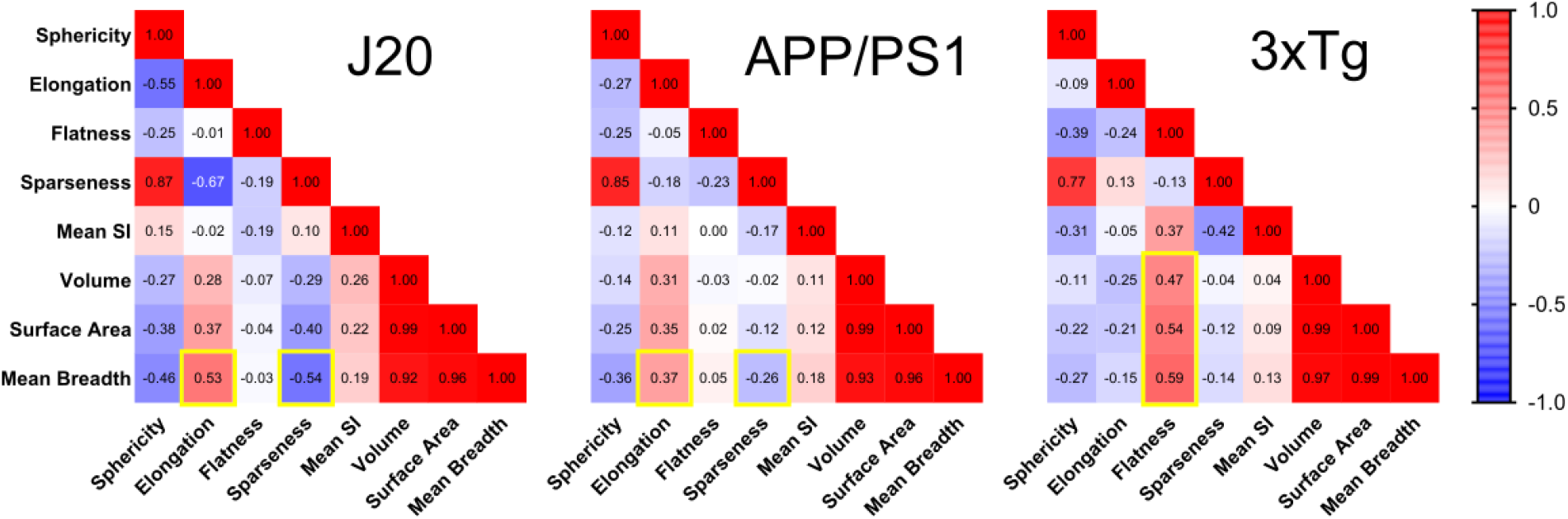
Pearson r correlation coefficients (color-coded from −1 = blue to 1 = red) between the 8 parameters.

While sphericity and sparseness were highly correlated in all three stains, flatness was correlated with surface area, volume and mean breadth in 3xTg only. In contrast, mean breadth in APP/PS1 and J20 was positively correlated with elongation, and negatively with sparseness.

## Discussion

The present study went beyond previous pioneer reports of Aβ detection with XPCT by:

i. studying a complete experimental group (10 brains in total vs 1-2 samples);
ii. comparing 3 different strains (vs a single one);
iii. and providing access to multiple morphometric parameters (while previous reports were confined to plaque number, volume and sphericity).

An additional contribution consisted in building and sharing a “biologist-friendly” analysis workflow, by assembling multiple Fiji plugins: segmentation editor, trainable WEKA segmentation 3D, MorpholibJ, 3D ImageJ Suite. We were especially interested in the ability of the workflow to correctly detect a significant proportion of Aβ plaques with very different appearances and morphologies. Hence versatility rather than completeness of the segmentation was the primary goal of the method. We were able to successfully segment numerous Aβ plaques, even when their refractive indices (and subsequent XPCT signals) were barely different from surrounding tissue, as in the case of APP/PS1. In future studies in a dedicated mouse strain, segmentation may be improved, by building a dedicated WEKA classifier combining a few slices from different animals, and/or by using several classes of objects to better discriminate Aβ plaques from neuronal tracts, and/or by using wild-type littermate brains to avoid false-positive detections and thus avoid human intervention.

Tested on three different transgenic mouse strains, the analysis workflow yielded an entirely new class of 3D parameters, with distribution easily measured on multiple plaques. We provide basic examples of how to handle this set of parameters: e.g., building correlation matrices (Fig. 6) or searching for spatial patterns ([Supplemental movie](https://box-ng.univ-lyon1.fr/s/X6gNsPY8bHKqZjb) showing the distribution of mean breadth values on APP/PS1 hippocampus). Hence, the present work leveraged 3D analysis to extract multiple morphological parameters of Aβ plaques which are hardly available with standard 2D histology or with serial microscopy with 3D reconstruction [24]. The development of such quantification pipelines is a necessary step for popularizing XPCT in neuroscience laboratories which need to phenotype transgenic animal cohorts. Moreover, this imaging modality could be instrumental in studying the prion-like properties of Aβ fibrillar aggregates. Typical experiments involve exogenously inoculated Aβ seeds (prion-like agents) which template and accelerate Aβ deposition in the host brain [25,26]. Hence non-destructive 3D imaging would be a great advantage in identifying the spreading routes of inoculated seeds [27]. Moreover, it was suggested that seed morphology influences the final morphology of disseminated Aβ deposits; but the existence of these “morphotypes” was inferred from 2D immunohistochemistry [28] and awaits 3D confirmation.

In this proof-of-concept report, 3xTg displayed a different type of Aβ plaque, with larger volume and area, greater elongation and flatness, and much higher signal intensity than J20 and APP/PS1. Aβ aggregation has been shown to start intracellularly in hippocampal neurons of the 3xTg strain, and it is interesting to note that neuronal bodies in the hippocampus showed high signal intensity in comparison to other strains (Fig. 1). Thus, the distinct aggregation cascade in this strain might translate into different extracellular morphotypes of Aβ plaque. While the mechanisms leading to these particularities remain elusive, these observations illustrate how XPCT can probe the pathophysiological microenvironment of brain tissue. In terms of morphometry, the study identified mean breadth as an original and discriminating parameter. Mean breadth is a quantity proportional to the integral of mean curvature over area (>0 for convex objects such as Aβ plaques). In this case, mean breadth can be defined as the average of the Feret diameter measured over a selected number of orientations. It correlated strongly with area and volume, but exhibited radically different correlation patterns with other parameters in 3xTg (Fig. 6). This shows that the analysis of 3D geometry can benefit from other parameters in addition to the often-reported sphericity and volume.

Though the pathophysiological relevance of Aβ shape remains unknown, the morphology of Aβ plaques gained interest with the recent development of brain clearing techniques [29–31]. In comparison, XPCT has the following advantages: i) ultra-fast sample preparation (around 1 hour, whereas 3D immunofluorescence typically requires several days’ incubation with antiamyloid anti-bodies), ii) limited and homogeneous variation in size through ethanol dehydration (whereas most clearing techniques drastically change brain size), and iii) whole-brain acquisition (whereas light-sheet imaging usually requires multiple acquisitions and fastidious stitching procedures). Therefore, XPCT can be positioned as a forefront technique, allowing high-throughput brain screening, and guidance for subsequent brain clearing and 3D immunohistochemistry on selected samples. XPCT is therefore unique as a whole-brain, label-free method for ex vivo amyloid imaging. The current limitations of XPCT are the following:

i. sensitivity of detection of Aβ plaques likely depends on size and location in the brain; this point could be addressed by more precisely adjusting tissue dehydration (by varying the percentage of ethanol) so as to get minimal anatomical contrast while preserving Aβ detectability;
ii. not all Aβ plaques seem detectable and the source of the XPCT contrast from Aβ plaques is not currently fully understood, though it is likely to arise from a local change in refractive index due to insoluble fibrillar Aβ [32], along with a possible contribution of endogenous metals entrapped in the plaque [33];
iii. the availability of synchrotron sources with XPCT capacity is restricted to 20-30 sites in the world; but several methods have been proposed to obtain phase-contrast images from a laboratory X-ray source [34–36].

In summary, we presented a complete workflow for ex vivo whole-brain imaging and quantification of Aβ pathology. Sample preparation was limited to reversible dehydration of tissue, which remained available for standard immunohistochemistry. Propagation-based XPCT produced unequalled image quality, with various concurrent types of anatomical information (white matter, vessels, neuronal organization). New 3D parameters, not attainable on routine immunohistochemistry, were successfully extracted from 3 transgenic Alzheimer models.

## Acknowledgments

This study was performed within the framework of LABEX PRIMES (ANR-11-LABX-0063) of Université de Lyon, within the “Investissements d’Avenir” program (ANR-11-IDEX-0007) operated by the French National Research Agency (ANR). The research was in part funded by the French ANR project NanoBrain (ANR15-CE18-0026). The study was supported by the European Synchrotron Research Facility (ESRF) by allocation of beam time (LS2292, MD1018, IN1041), and the authors would like to thank ESRF local contact Lukas Helfen. Carlie Boisvert was supported by Mitacs Globalink Canada (travel grant). 3xTg and WT brains were kindly provided by Dr Catherine Lawrence of the University of Manchester. Paraffin embedding and microtome slicing were performed at the CIQLE Imaging Platform, University of Lyon, with the help of Annabelle Bouchardon and Batoule Smatti.

## Supplemental Figures

**Figure S1-.**
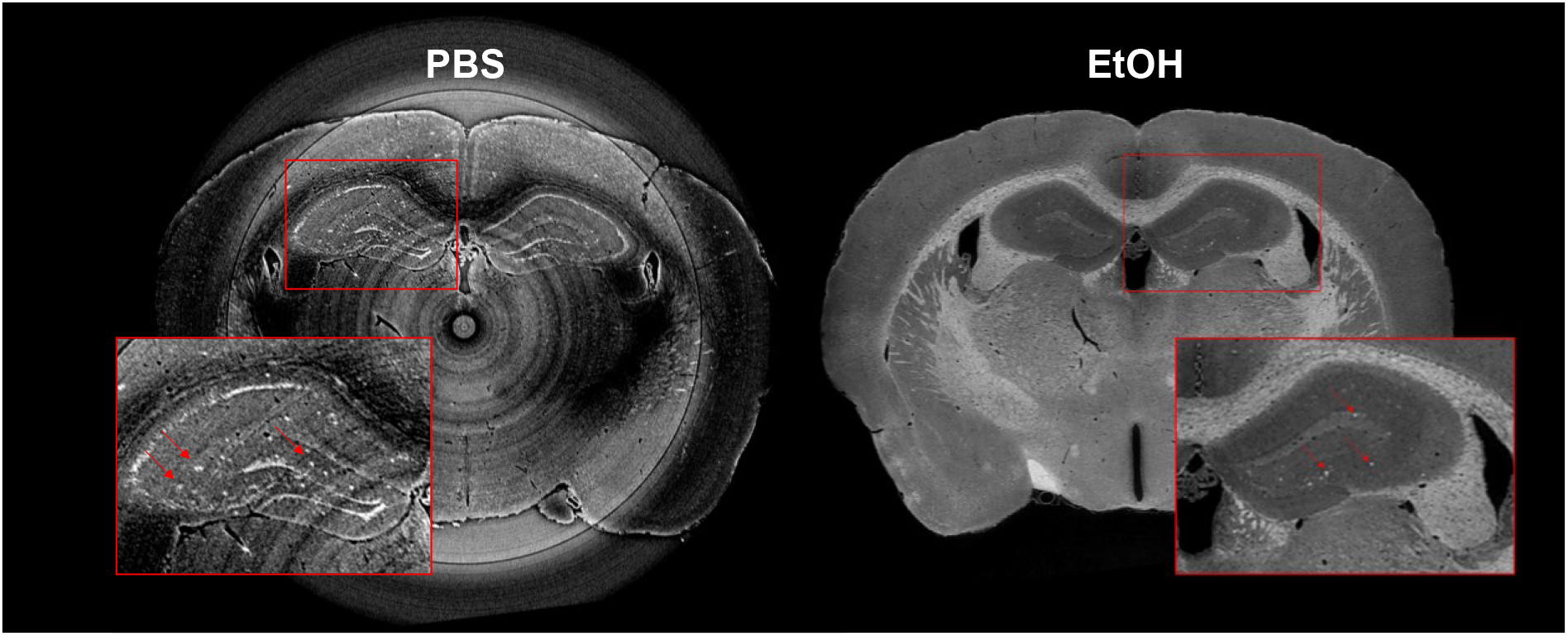
Single-slice XPCT images from J20 brains with (right) and without (left) dehydration in ethanol.

**Figure S2.**
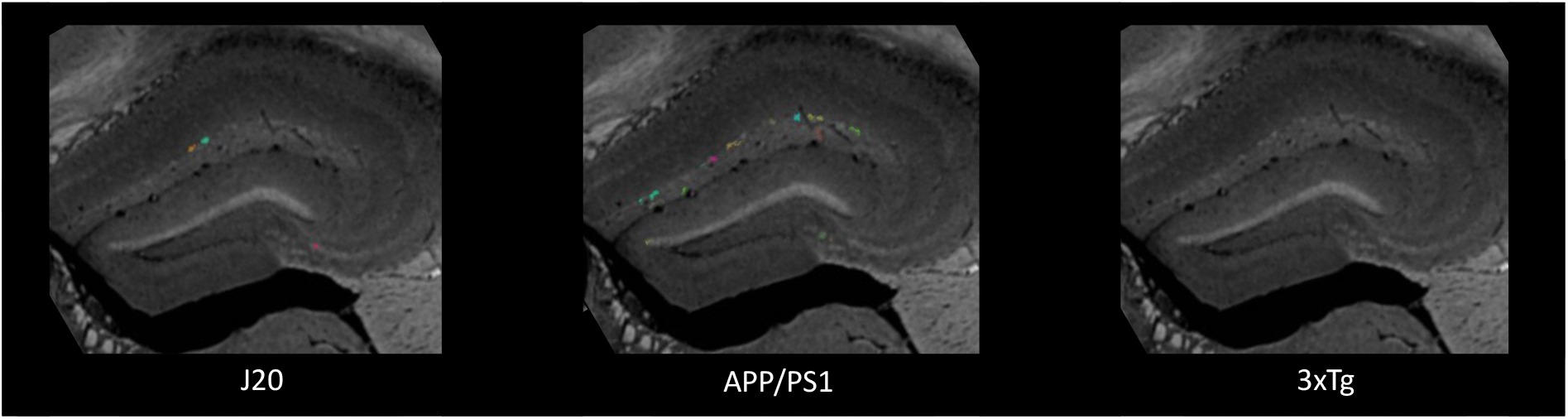
Results of trainable WEKA segmentation using a strain-specific classifier applied on the same non-transgenic C57/Bl6 brain. Colored labels represent false positive detections of plaque-like signals.

**Figure S3.**
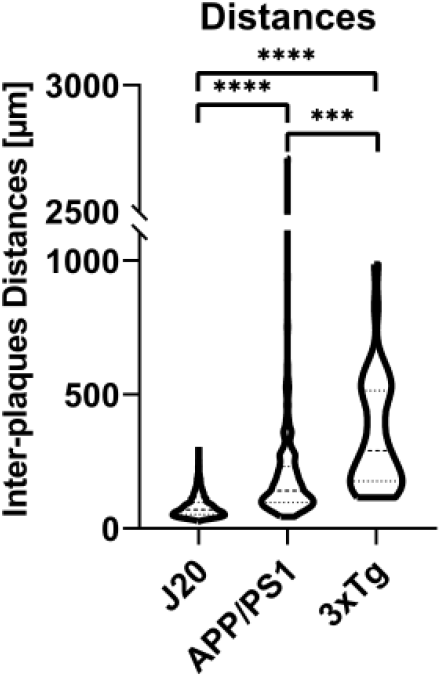
Violin plots of nearest-neighbour distances extracted from the population of segmented Aβ plaques. The present results are reported to highlight feasibility but may be biased because only a subset of individual plaques was selected in APP/PS1, and closest pairs of J20 plaques couldn’t be resolved.

